# HP1γ binding pre-mRNA at intronic repeats increases splicing fidelity and regulates alternative exon usage

**DOI:** 10.1101/686790

**Authors:** Christophe Rachez, Rachel Legendre, Mickaël Costallat, Hugo Varet, Jia Yi, Etienne Kornobis, Christian Muchardt

## Abstract

HP1 proteins are best known as markers of heterochromatin and gene silencing. Yet, they are also RNA-binding proteins and the HP1γ/Cbx3 family member is present on transcribed genes together with RNA polymerase II, where it regulates co-transcriptional processes such as alternative splicing. To gain insight in the role of the RNA binding activity of HP1γ in transcriptionally active chromatin, we have captured and analyzed RNAs associated with this protein. We find that HP1γ specifically recognizes hexameric RNA motifs and coincidentally transposable elements of the SINE family. As these elements are abundant in introns, while essentially absent from exons, the HP1γ RNA binding activity tethers unspliced pre-mRNA to chromatin via the intronic region and limits the usage of intronic cryptic splice sites. Thus, our data unveil novel determinants in the relationship between chromatin and co-transcriptional splicing.

## INTRODUCTION

Maintenance and propagation of the transcriptionally inactive heterochromatin extensively relies on Heterochromatin Protein 1 (HP1), a family of proteins identified in a large variety of species, ranging from fission yeast to man^1^. Mammals typically express three isoforms of HP1, namely HP1α/CBX5, HP1β/CBX1, and HP1γ/CBX3, each with unique subnuclear localization patterns^2, 3^. HP1 proteins bind histone H3 trimethylated at Lysines 9 (H3K9me3) via their N-terminal chromodomain^4, 5^. At their C-terminus, a chromoshadow domain ensures dimerization and mediates interaction with numerous molecular partners characterized by the presence of a PXVXL motif^6^. In-between these two structurally very similar globular domains, an unstructured region known as the Hinge harbors both DNA and RNA binding activity^7^.

The RNA binding activity of HP1 proteins seems very important for their molecular function. In the fission yeast *Schizosaccharomyces pombe*, HP1/Swi6 associates with noncoding transcripts expressed in centromeric heterochromatin, and its silencing activity relies on a mechanism involving RNAi-dependent degradation of these transcripts^8^. This process was later shown to involve a dynamic trafficking of HP1/Swi6 between its free, H3K9me-bound and RNA-bound forms, leading to the repression of heterochromatin^9^. For murine and human HP1*α*, the RNA binding activity of the Hinge is essential for the targeting of this protein to heterochromatin, possibly even more so than the H3K9me3 histone modification^10–12^. While HP1 proteins may bind multiple families of RNA species^13^, mouse HP1*α* was shown to specifically bind pericentromeric RNA transcripts from the major satellites, a family of repeats particularly abundant in pericentromeric heterochromatin^11, 14^.

Beyond their role in structuring heterochromatin, HP1 proteins also functions as regulators of euchromatic transcription. For instance, at the promoters of many inducible genes involved in development or in cellular defense, they participate in the transient silencing of transcription while awaiting stimulation^15–20^. But HP1 proteins, in particular HP1γ in mammals, are also present inside the coding region of genes^21^, a localization which is not always correlated with H3K9me3 ^19^. The association of HP1γ with transcribed genes is consistent with a role for this protein in co-transcriptional mechanisms such as termination^22^, and regulation of alternative splicing^23–27^.

Splicing is a maturation process of RNApol2 transcripts catalyzed by the Spliceosome complex, and leading to the formation of mature mRNA by excision of introns and joining of exons. Most human genes undergo alternative splicing which gives rise to multiple mRNAs from a single gene locus^28, 29^. As splicing is mostly co-transcriptional and occurs in the close vicinity of chromatin, it is influenced by a large number of chromatin-associated factors^30, 31^. In this context, we have shown earlier that recruitment of AGO proteins and HP1γ to CD44 and other genes favors intragenic H3K9 methylation and affects the outcome of alternative splicing by targeting the spliceosome to specific sites inside the gene body^25^.

Our study on the CD44 gene also unveiled an interaction between intragenic chromatin and pre-mRNA which was dependent on HP1γ and seemed to modulate the outcome of splicing^25^. To gain further understanding of this HP1-dependent relationship between chromatin and transcripts, we have here analyzed the genome-wide association of HP1γ with RNA by a chromatin-enriched RNA immunoprecipitation (RNAchIP) assay. We find that HP1γ preferentially associates with intronic regions, due to the presence therein of hexameric motifs. Consequently, HP1γ-bound RNAs are also enriched in B4 SINEs, a family of euchromatic transposable repeat elements which harbors high proportions of these hexameric motifs. The consequence of this RNA binding by HP1γ is a tethering of unspliced pre-mRNA to chromatin via the intronic region. This way, HP1γ limits the usage of intronic cryptic splice sites. These observations reconcile the heterochromatic and euchromatic functions of HP1, by showing that its role in mRNA maturation, alike its role in heterochromatin structuring, relies on its ability to associate with repeat-encoded RNAs.

## RESULTS

### HP1γ associates with chromatin-enriched RNA

To better understand the relationship between HP1γ and RNA, beyond its classical role in heterochromatin formation, we used a genome-wide approach to assay the interactions between HP1γ and RNA on chromatin (RNAchIP; Fig. 1a). For this, we used a modification of our previously described strategy to solubilize native chromatin and produce chromatin-enriched RNA fragments suitable for immunoprecipitation^25^ (Supplementary Fig. 1d). HP1γ -/- (KO cells) mouse embryonic fibroblast (MEF)-derived cell lines, re-complemented with FLAG-tagged HP1γ (HP1γ cells) as previously described were used in these assays^16^. These cells expressed ectopic FLAG-tagged HP1γ at the same level as the endogenous HP1γ in WT MEFs (Supplementary Fig. 1a). As in our previous studies, the cells were treated or not with the phorbol ester PMA, an activator of the PKC signaling pathway, in order to observe the cellular response to stress on chromatin and transcription. Nuclei were isolated to obtain a chromatin-enriched RNA fraction, (Supplementary Fig. 1a and b). Our procedure was modified to include a limited cross-linking step in order to stabilize association of HP1γ with RNA, together with stringent immunoprecipitation conditions to favor direct protein-RNA associations. Under these conditions, anti-FLAG antibody did not precipitate any of the HP1γ-interacting proteins we tested (HP1*α*, H3, RNA polymerase II) (Supplementary Fig. 1b). When quantified by RT-qPCR, HP1γ was found to associate with chromatin-enriched RNA from the *Fosl1* locus with a 20-to 25-fold enrichment compared to KO cells. IP in KO cells consistently showed very low to undetectable RNA levels (Supplementary Fig. 1c). Genome-wide analysis of the RNAchIP by HP1γ (IP) versus RNA detected in the chromatin fraction (input) was then performed using Illumina sequencing on biological triplicates. Reads were then mapped onto the mouse genome. In a first approach, RNAchIPseq data were computed by a transcriptome analysis. Normalized read counts per gene body (input and IP) revealed that RNA enrichment in IP is correlated with the levels of chromatin-associated RNA (Fig. 1c). In addition, stimulation of cells with PMA increased both input and IP read density on stress-responsive genes such as the *Fosl1* gene (Fig. 1b middle). Together, these readings indicated a positive correlation between production of transcripts and their binding to HP1γ whether or not transcription was stimulated with PMA (Fig. 1b, c), in agreement with earlier observations correlating HP1γ recruitment on DNA with gene expression^26^. However, while several genes were highly enriched in IP (see *Gcnt4* or *Fosl1*, Fig. 1b), transcripts from several other genes including most histone genes were depleted (*Hist1h4a* and *Hist1h3a*, Fig. 1b right), indicating that HP1γ does not equally associate to all transcripts.

**Fig. 1.**
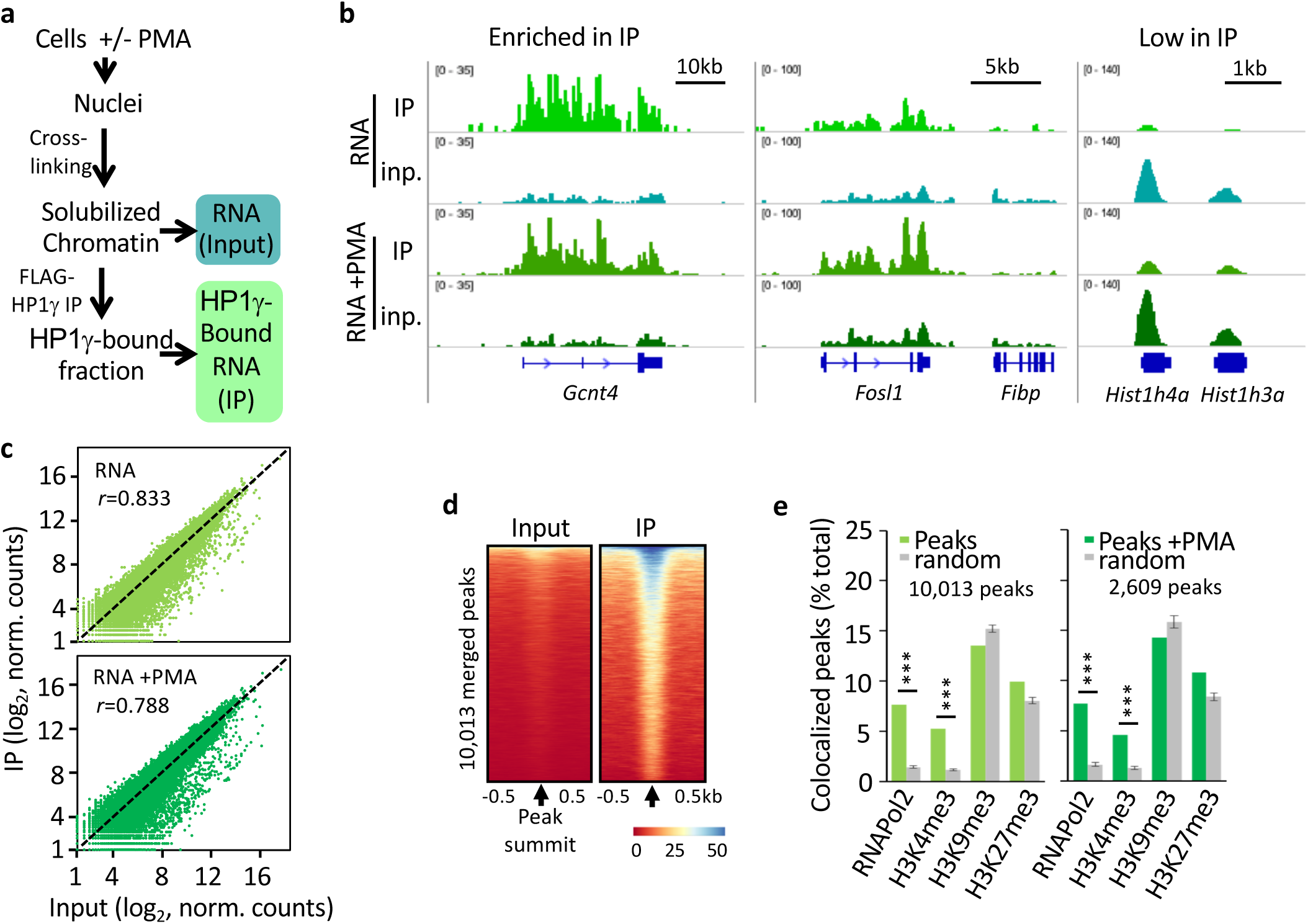
HP1γ associates with introns of pre-messenger RNA on chromatin. **a,** Scheme of the strategy used to assay HP1γ association with RNA on chromatin. **b**, Genome views of RNA read density profiles on representative genes enriched (left) or low (right) in HP1γ RNAchIP (IP) and in input RNA (inp.) from HP1γ cells stimulated or not with PMA. **c**, Genome-wide scatter plot of IP and input normalized (norm.) RNA read counts per gene for a representative sample of the triplicates; n=21,754; *r*, Pearson’s correlation coefficient between IP and input. **d**, Heat maps of RNA IP and input signal centered on the summit of RNAchIP peaks of IP versus input in unstimulated samples detected by MACS2 analysis (10,013 peaks, merged between triplicates). **e**, Percentage of RNAchIP peaks colocalized with indicated chromatin features in MEF samples from ENCODE database (green bars). Colocalization was evaluated by comparison with the list of merged peaks whose genomic location was randomized among genes (gray bars). Graphes represent mean and s.d. for the randomized peaks, *n*=100. P-values indicate significantly higher difference between two conditions (* P < 0.01, ** P < 0.005, ***P < 0.001; two-tailed Student’s *t*-test).

Within individual genes, the distribution of HP1γ-associated RNA appeared to be fluctuating along the gene body (Fig. 1b). To detect regions of local RNA enrichment by IP, we searched for peaks of IP versus input RNA in all replicates. By merging peaks conserved in at least two of the three replicates, we identified 10,013 and 2,603 peaks, in unstimulated and in PMA-stimulated samples, respectively (Fig. 1d,e). As HP1γ chromatin localization may rely on determinants other than the H3K9me3 histone mark, we looked at the chromatin context of the peaks. For this purpose, we screened for overlaps between these peaks and several chromatin features obtained by ChIP in MEF cells and available in the ENCODE database, compared to random. Interestingly, the peaks colocalized best with RNA polymerase II and H3K4me3, two features associated with active transcription. No significant colocalization was observed with H3K9me3 and H3K27me3, the two major histone modifications associated with heterochromatin (Fig. 1e). These data clearly linked RNA-associated HP1γ with sites of active transcription.

### HP1γ-associated RNA is enriched in CACACA and GAGAGA motifs

We next investigated whether sequence specific motifs could be found within the peaks. Peak-motif analysis using RSAT on stranded sequences of all RNA peaks revealed an enrichment in CACACA motifs (e-val. 4.8 e-88), and to a lesser extend in GAGAGA motifs (e-val. 1.8 e-10) in more than half of our peaks (57%; Fig. 2a). Consistent with an enrichment in CACACA sequences in the RNA co-immunoprecipitating with HP1γ, we observed an accumulation of reads around all intragenic CACACA motifs at expressed genes, visualized by an increased average distribution of the reads from IP compared to input RNA around these motifs (Fig. 2b), and the clustering of reads at a large number of intragenic loci centered on CACACA motifs (Supplementary Fig. 2a). Noticeably, we did not find any enrichment in the TGTGTG motif, antisense to the CACACA motif (Fig. 2b).

**Fig. 2.**
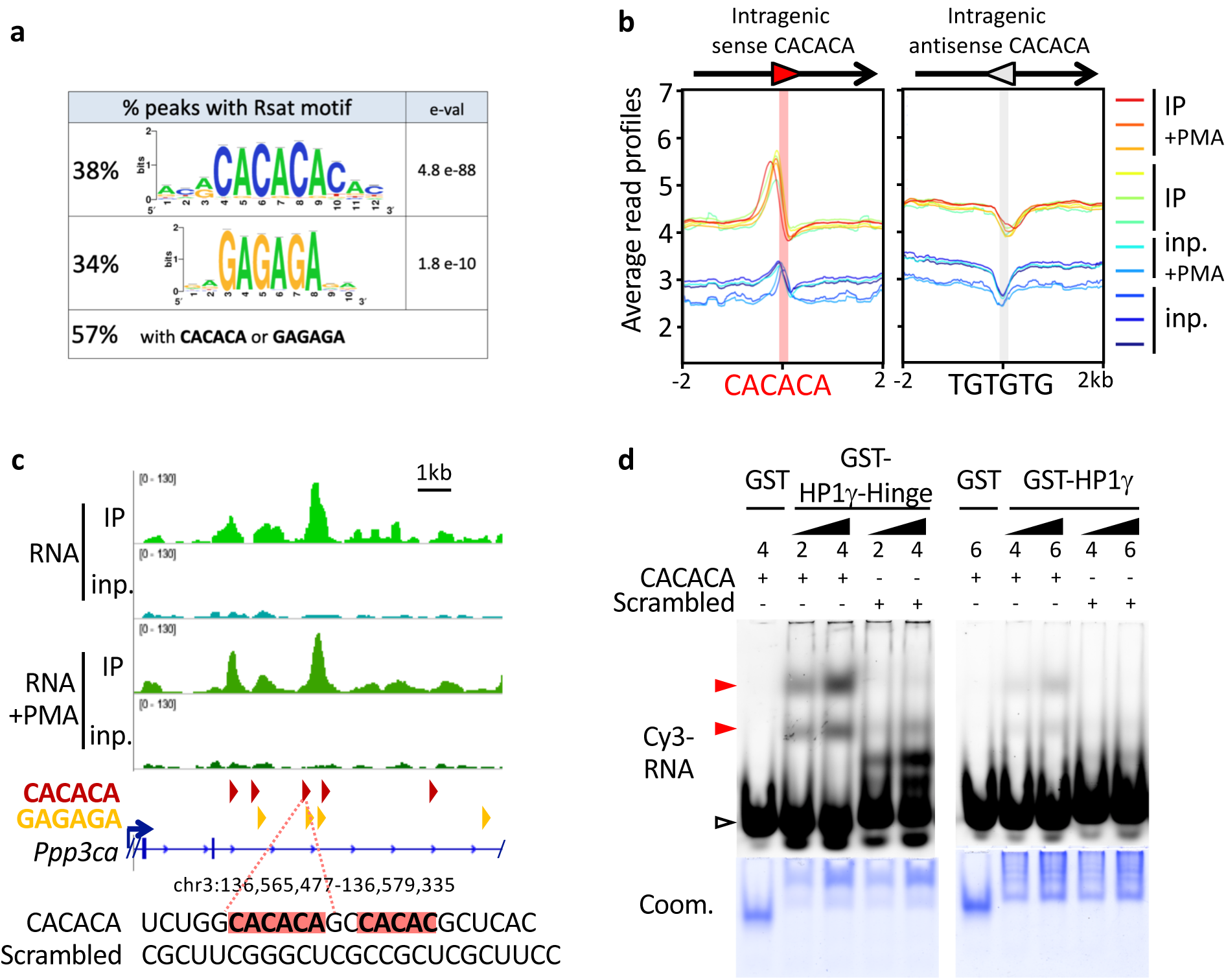
HP1γ-associated RNA is enriched in CACACA and GAGAGA motifs. **a**, Consensus motifs discovered among RNAchIP peaks with the RSAT pipeline, together with the percentage of peaks containing at least one exact hexameric CACACA or GAGAGA motifs or both. e-val. represents the expected number of patterns which would be returned at random for a given probability. **b**, Distribution profiles of average RNAchIP signal in both IP (warm colors) and input (blue colors) over ±2kb centered on intragenic CACACA hexameric motifs oriented in the same orientation (sense) or in opposite orientation (antisense) relative to the overlapping annotated transcript. The antisense CACACA motifs were obtained by querying the transcript sequences with the TGTGTG motif. **c**, Top, representative example of a locus on the *Ppp3ca* gene surrounding an RNAchIP merged peak, showing the RNAchIP read density as in Fig. 1b. Red and orange arrowheads correspond to oriented CACACA and GAGAGA motifs, respectively. Bottom, sequence of the RNA surrounding a CACACA motif highlighted in red, as well as a second a neighboring imperfect motif. The sequence was used to design a CACACA-containing RNA oligonucleotide probe (CACACA), compared to a control probe (scrambled) devoid of any related motif. **d**, Gel mobility shift assay of bacterially expressed, purified GST proteins fused to HP1γ or HP1γ-Hinge domain, and tested for their direct interaction with the Cy3-labeled RNA oligonucleotide probes depicted in **c**. RNA probes in the gels were detected by their Cy3 fluorescence. Total GST-fusion protein loading was visualized by Coomassie blue staining (Coom.). Representative of 3 independent experiments.

To test whether the contact between HP1γ and the RNA was direct, we identified a representative RNA peak within the Ppp3ca gene which overlaps both CACACA and GAGAGA motifs (Fig. 2c), then used the sequence overlapping one of these motifs *in vitro* as an RNA probe in gel mobility shift assays. This probe was compared to a scrambled RNA devoid of any consensus motif (Fig. 2c,d). Purified GST-HP1γ expressed in bacteria (Supplementary Fig. 2b) demonstrated a clear specific affinity for the CACACA motif. Since earlier studies have shown that an isolated hinge domain in HP1 proteins has more DNA- and RNA-binding activity *in vitro* than the full-length proteins^10, 12^, we also included in this experiment a fragment of HP1γ corresponding to the Hinge region (covering amino acids 67 to 101, Supplementary fig. 2b). This isolated HP1γ domain showed a clear binding to the CACACA-containing probe while showing no affinity for the control scrambled RNA (Fig. 2d). Similarly, purified HP1γ-hinge was found to directly bind an RNA probe containing the GAGAGA motif (Supplementary Fig. 2c). HP1γ has therefore the capacity of directly interacting with RNA in a sequence specific manner, suggesting that the RNA enrichment seen in IP is due to direct HP1γ/RNA associations at specific positions enriched in CACACA or GAGAGA motifs.

### HP1γ-associated RNA is enriched in SINE repeat motifs

Heterochromatin-based silencing mechanisms may occur within active chromatin on repeated sequences such as interspersed repeats (SINEs, LINEs) or endogenous retroviruses (LTRs)^32, 33^. We therefore asked whether HP1γ-associated RNA could be enriched over different classes of repeated elements annotated in the RepeatMasker database. When aligned onto repeats with our multimapping alignment procedure, sequencing reads showed best enrichment on LTR and SINE repeats (Fig. 3a).

**Fig. 3.**
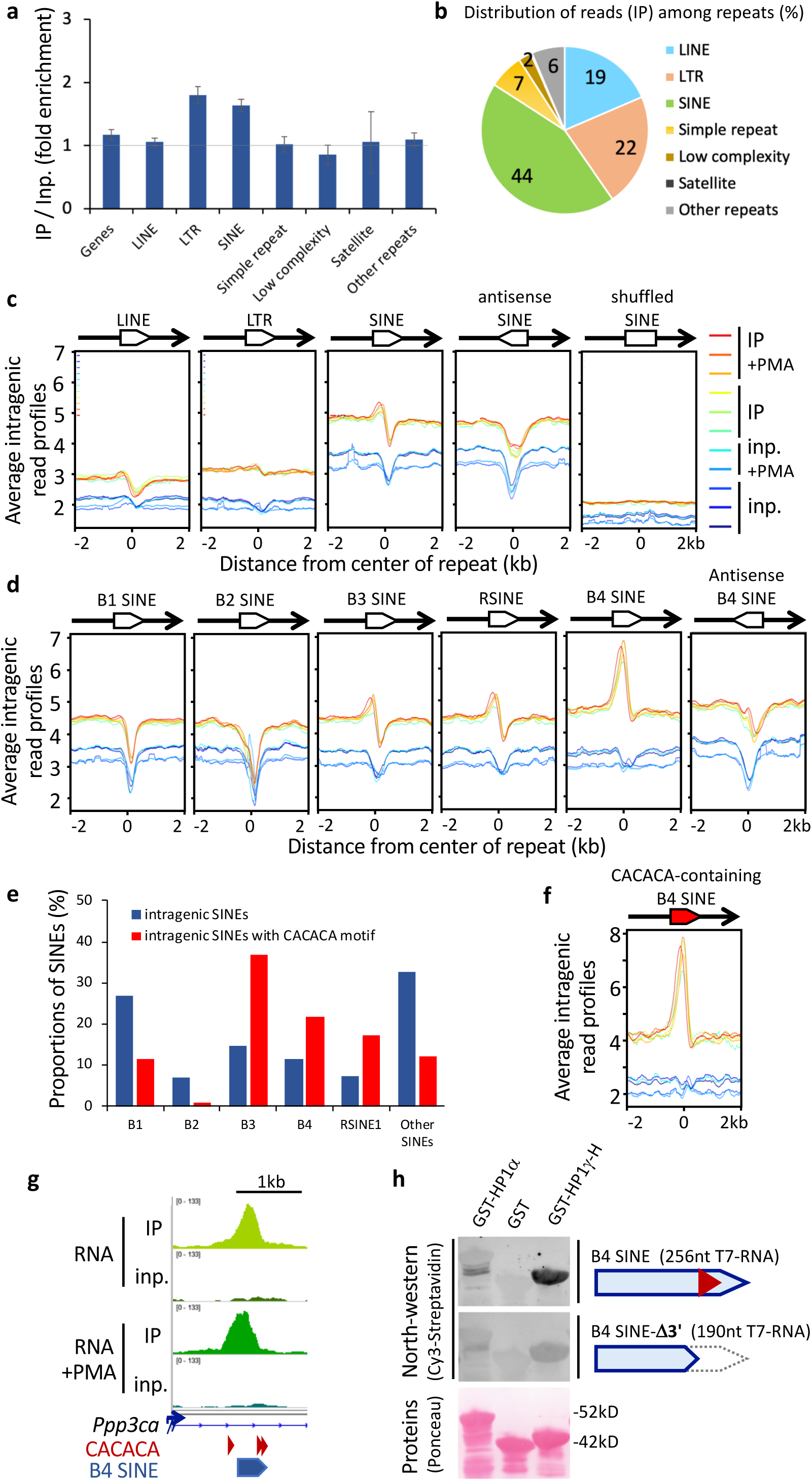
HP1γ-associated RNA is enriched in oriented SINE repeat motifs. **a,** Fold enrichment values on all repeat masker classes and on Refseq genes, on the basis of all uniquely aligned reads, shown as mean and s.d., *n=*6 biological replicates. **b,** Pie chart of the distribution of RNA sequencing reads in IP per repeat masker classes counted as in **a**, as a percentage of all repeats. **c**, Profiles of average RNAchIP signal as in Fig. 2b over ±2kb centered on intragenic LINE, LTR, or SINE repeats annotated in Repeat masker database. All repeats are oriented in the same orientation (sense) relative to the overlapping annotated transcript, unless otherwise specified (antisense, shuffled), and as depicted by an oriented white box overlapping the transcript in 5’ to 3’ orientation (black line with arrowhead). **d**, Profiles of average RNAchIP signal, as in **c** centered on the 5 major families of intragenic SINE repeats. **e**, Respective proportions of the families of both SINE and CACACA-containing SINE repeats, as a percentage of total intragenic SINEs. **f**, Profiles of average RNAchIP signal, as in **d** centered on the subset of intragenic B4 SINEs containing a consensus CACACA motif. **g**, Genomic position of a B4 SINE repeat and CACACA consensus motifs at the location of the RNAchIP peak depicted in Fig. 2c. **h**, North-western blot assay showing direct association between the indicated bacterially expressed, purified GST-fusion proteins and *in vitro* transcribed, biotinylated RNA probes based on the sequence of the B4 SINE depicted in **g**. Top panels, binding of a 256nt probe corresponding to the CACACA-containing B4 SINE sequence, was compared to an identical B4 SINE deleted of its CACACA by truncation of its 3’ portion (B4 SINE-Δ3’). RNA probes hybridized on the membranes were detected by their Cy3 fluorescence. Total GST-fusion protein loading was visualized by Ponceau S staining. Representative of two independent experiments.

We then focused on SINEs, LINEs, and LTRs which are the most abundant repeats within gene bodies (Fig. 3b, Supplementary Fig. 3a). Profile plots for IP read counts over these repeats confirmed that the best enrichments were to be found in the neighborhood of SINEs (Fig. 3c – note that the body of the various repeats are plotted as valleys because multimapping reads have been filtered away). Enrichment was most obvious on the B4 SINE family, and also more pronounced on the B3 and RSINE families, while undetected on the B1 and B2 SINEs (Fig. 3d). Consistent with a preference of HP1γ for CACACA sequences, B3, B4 and RSINEs contain this hexamer motif in their sequence more frequently than other SINEs (Fig. 3e). In addition, average profiles for IP read counts over CACACA-containing B4 SINEs showed a better enrichment than that observed over B4 SINEs in general (Fig. 3f, compare with Fig. 3d B4 SINE panel). Noticeably, only SINEs in the same orientation as their host gene were found enriched (Fig. 3c, d). Consistent with this, the reverse-complement sequence of the B4 SINEs were not enriched in CACACA motifs (Supplementary Fig. 3b).

Finally, we tested a SINE sequence for direct binding of HP1γ. As in Fig. 2c, we selected a sequence corresponding to the IP peak present inside the Ppp3ca gene, which happens to contain a B4 SINE (Fig. 3g). Because of the size of the probe (256 nt), a Northwestern assay was preferred. This assay confirmed that an RNA probe designed around the SINE sequence had the capacity of establishing direct contacts with HP1γ (Fig. 3h). This association was dependent on the 3’ region of the probe which contains the CACACA motifs (Fig. 3h). Altogether, our results are consistent with a binding of HP1γ to SINE-containing pre-messenger transcript, not to individual SINE transcript, and suggest that the intragenic SINE repeats, concomitantly to the presence of a CACACA motif, constitute a targeting motif for the association of HP1γ with chromatin-enriched transcripts.

### The CACACA motif affects HP1γ association with introns of pre-messenger transcripts on chromatin

Transcripts from most histone genes were depleted from the IP fraction as compared to the input in the HP1γ RNAchIP (*Hist1h4a* and *Hist1h3a*, Fig. 1b). Since most histone genes are intron-less, we monitored IP RNA enrichment on genes sorted according to the number of exons they contained (Fig. 4a, Supplementary Fig. 4). Quantification of reads over transcripts of intron-less genes appeared depleted in IP (Fig. 4a, orange box plot, 1exon), while all intron-containing genes were enriched in IP in a similar manner, independently of the number of exons (Fig. 4a, blue box plots, 2, 3, and 4 exons). Further quantification of reads in IP showed a 1.4-fold enrichment over introns and a depletion (0.7-fold) over exons when compared to input (Fig. 4b). Finally, we quantified in IP the reads aligning on junctions between two exons and therefore functioning as reporters of completed splicing reactions. The proportions of such exon-exon junction reads were significantly lower in IP relative to input (Fig. 4c). Together, these observations indicate that HP1γ associates preferentially with intronic regions. Since most genes are spliced co-transcriptionally, our results further suggest that HP1γ associates preferentially with pre-mRNAs prior to maturation by splicing.

**Fig. 4.**
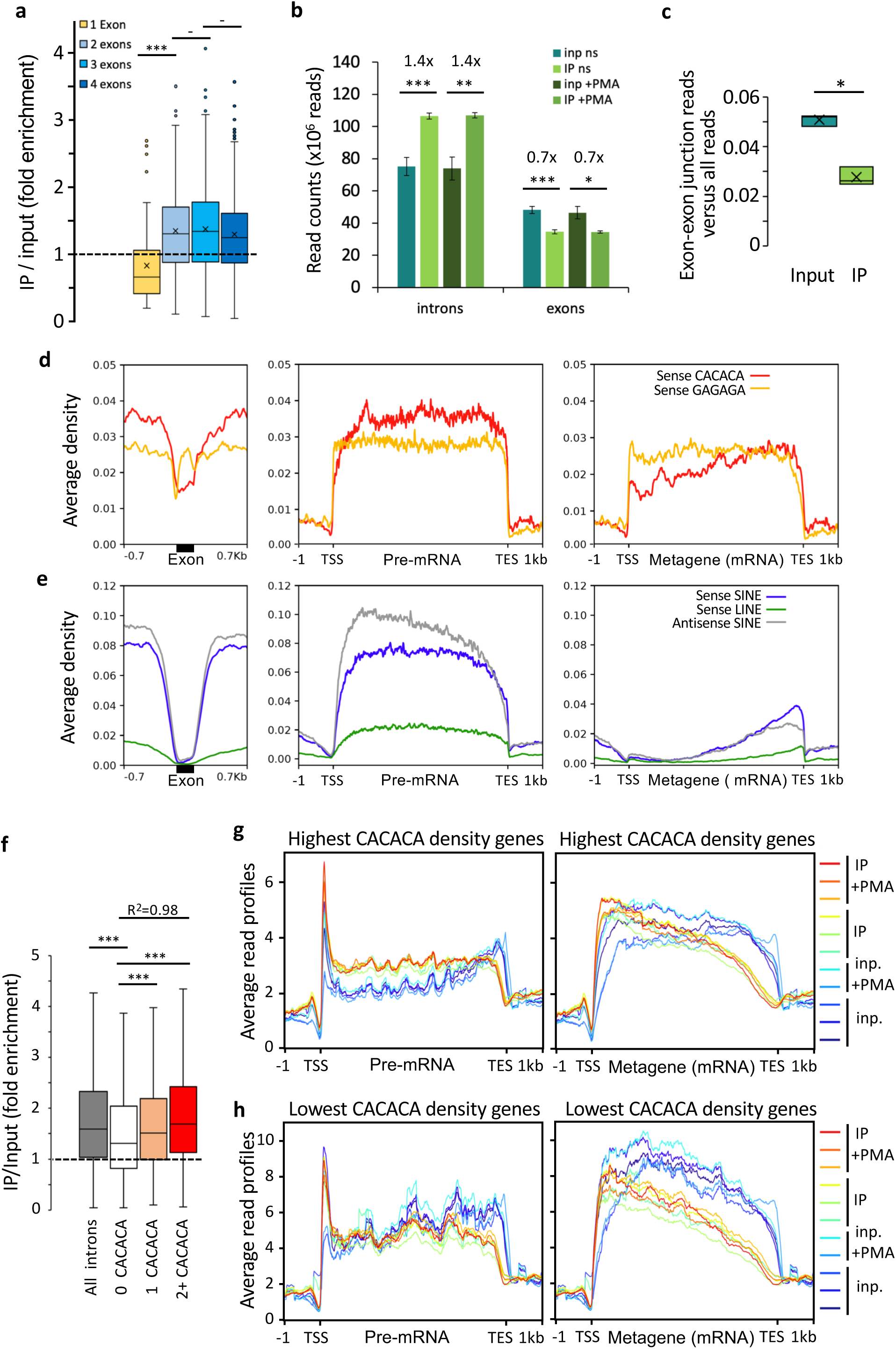
The density of CACACA motifs affects HP1γ association with transcripts on chromatin. **a**, Box plot showing the distribution of fold enrichment values per genes sorted by exon number, *n=* 66, 247, 389, and 574 genes with 1, 2, 3, and 4 exons, respectively. All box boundaries in Fig.4 represent 25th and 75th percentiles; center line represents median; whiskers indicate ±1.5×IQR; points are values of outliers. All P-values in Fig.4 are * P < 0.01, ** P < 0.005, ***P < 0.001; two-tailed Student’s *t*-test. Box boundaries represent 25th and 75th percentiles; center line represents median; whiskers indicate ±1.5×IQR; points are values of outliers. **b,** Reads from input and IP counted in unstimulated and PMA stimulated triplicates give significant enrichment values of 1.4 and 0.7 on introns and exons, respectively, shown as mean and s.d., *n*=3 biological replicates. **c**, Box plot of reads belonging to spliced transcripts versus all reads in all six input or RNAchIP samples calculated as a ratio between split reads and total reads; x represents average, *n*=6 biological replicates. **d, e,** Average distribution profiles of the depicted hexameric motifs (**d**), or SINE and LINE repeats (**e**) on RefSeq genes, computed over exons and on ±0.7kb of surrounding introns (left panels), or over entire gene bodies including introns (pre-mRNA; middle panels), or as metagenes over exons only (Metagene mRNA; right panels). **f,** Box plot as described in **a** showing the distribution of fold enrichment values on series of introns of comparable size (4kb in average, ranging from 3 to 5kb) sorted by number of sense CACACA motifs per intron (2+, two or more motifs), *n=* 5782, 899, 960, and 3234 introns for all introns, 0, 1, and 2+ CACACA categories, respectively. R^2^, Linear regression coefficient between 0, 1, and 2+. **g, h,** Average read profiles as in Fig. 2b computed as in **d** and **e** on two groups of RefSeq genes, in the size-range 4 to 15kb, containing the highest (**g**) or lowest (**h**) CACACA motif density.

To explore whether binding of HP1γ to introns could be explained by the sequence specificity of this protein, we profiled the distribution of the HP1γ-targeting hexamer motifs within exons and in their surrounding introns (Fig. 4d,e). This approach uncovered that the GAGAGA motif is excluded from the exon boundaries likely due to the strict sequence constraints linked to the definition of these boundaries. The profile of the CACACA motif was different and revealed a clear depletion of this motif from exons (red profile). Consistent with this motif being predominantly located in introns, pre-mRNAs were found to contain higher density of CACACA motifs than their mature mRNA counterparts, devoid of introns (Fig. 4d, compare red profiles in middle and right panels). An even stronger intronic enrichment was seen when considering intragenic SINE repeats (Fig. 4e, left, blue and grey profiles). Indeed, unlike LINEs (green profile), SINEs were essentially absent from exons, but present inside introns throughout the gene body (compare metagene versus meta cDNA profiles, Fig. 4e).

To probe for a correlation between the CACACA content of introns and their association with HP1γ, we selected introns of comparable size (4kb in average, ranging from 3 to 5kb), and sorted them according to their CACACA content. This approach revealed a significant increase in IP over input ratio when the number of CACACA motifs increased within introns (compare categories of introns containing no motif versus 1, and 2 or more motifs, Fig.4f). To address the impact of the CACACA motif at the scale of entire genes, we sorted genes into two categories according to their global CACACA density (1700 genes above 0.5 hexamers/kb and 900 genes below 0.3 hexamers/kb, Fig. 4g and h, respectively, genes were 9kb in average, ranging from 4 to 15kb). These two categories clearly showed opposite profiles of average read density in IP. The genes containing the highest CACACA density were enriched in IP while the genes having the lowest density were depleted, affecting both pre-messenger RNAs (compare relative position of red IP profiles and blue input profiles in Fig. 4g and h, left panels), and mature mRNAs on chromatin (compare Fig. 4g and h, right panels). These profiles highlight a positive correlation between the density in CACACA motifs and the enrichment in IP over whole genes, and suggest that this global enrichment results from the multiplicity of discrete HP1γ binding motifs. For both categories of genes, the relative position of the average IP and input profiles were also shifted when considering metagenes rather than pre-mRNAs profiles (compare left and right panels Fig. 4g, h). This clearly suggests that mature transcripts are released from HP1γ, and it is consistent with the preferential association of HP1γ with introns that we described above. Interestingly, the genes with the highest CACACA motif density show globally a lower average read density than the lowest CACACA category (compare Fig. 4g and h). From this inverse correlation, we hypothesize that HP1γ may have a positive impact on the maturation of these transcripts and their release from chromatin in accordance with previous studies ^26^.

### HP1γ has an impact on the fidelity of RNA splicing which is correlated with intronic RNA binding

We have previously demonstrated a role for HP1γ in the regulation of alternative splicing^25^. To investigate a possible link between the binding of HP1γ at intronic repeats and its impact on splicing, we performed a transcriptome analysis on the HP1γ-expressing (HP1γ) and HP1γ-null (KO) cells lines (Supplementary Fig. 5a). The differences in splicing decisions between the two cell lines were quantified with the MAJIQ algorithm^34^. As anticipated from earlier studies, this approach led us to identify a number of splice junctions affected by the presence of HP1γ (178 high confidence differential splicing events in total, see example Fig. 5a, Pie chart in Supplementary Fig. 5b, and list of events in Supplementary Table 1). However, we noted that in approximately one third of these differential splicing events (64 events), the loss of HP1γ was correlated with increased usage of cryptic splice sites involving intronic sequences (see example Fig. 5b and categories of splicing events in Supplementary Fig. 5b). This observation suggested a role for HP1γ in promoting usage of genuine splice sites over cryptic intronic ones. To further document this, we evaluated the average length of excised introns in the two cell lines. While genuine junctions were unaffected, junctions to cryptic splice sites were significantly shorter in the absence of HP1γ (pVal=2×10^-7^), strongly suggesting that additional cryptic splice sites become available to the spliceosome when this protein is missing (Fig 5c).

**Fig. 5.**
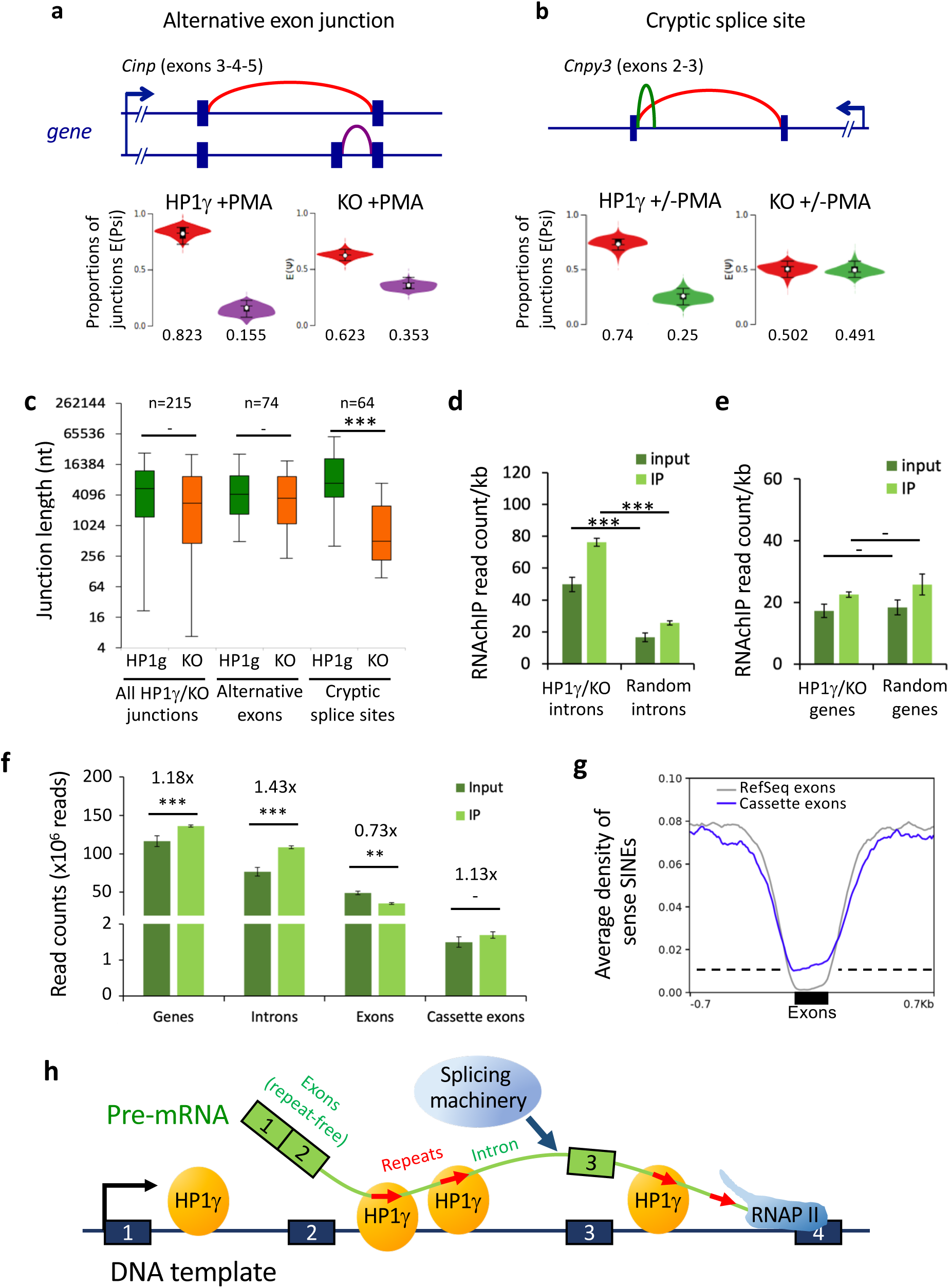
Intronic RNA binding by HP1γ has an impact on RNA splicing. **a,b**, HP1γ-dependent differential splicing events (junctions) detected by MAJIQ between transcriptomes of HP1γ and KO cells. Top, genome views showing annotated exon-exon junctions (red, panels **a** and **b),** an annotated alternative splicing event (purple, panel **a**), or a *de novo* cryptic splicing event increased in KO (green, panel **b**). Below, violin plots of proportions of the different junctions detected above in HP1γ and KO transcriptomes. **c**, Box plot, as described in Fig. 4a, depicting the length of exon junctions found differentially regulated by MAJIQ between HP1γ (green) and KO (orange) in three categories, all differential HP1γ/KO junctions, annotated alternative exon junctions, non-annotated *de novo* cryptic splice sites detected in KO only; n, differential events. **d**, RNAchIP read density per kb on the indicated introns for input and IP in combined samples from both untreated and PMA stimulated cells. HP1γ/KO introns are all intronic intervals (n=272) between exon junctions differentially detected in HP1γ and KO. Random introns are a matching library of random intronic intervals. **e**, RNAchIP read density per kb on the genes corresponding to the introns tested in **d**. **f**, Reads from input and IP counted on the indicated features in combined unstimulated and PMA stimulated triplicate samples. All graphes represent mean and s.d. (** P < 0.005, ***P < 0.001; two-tailed Student’s *t*-test). **g**, Average distribution profiles of sense SINE repeats on indicated exons and on ±0.7kb of surrounding introns. **h**, Model of HP1γ-dependent impact on splicing events via its binding to repeated intronic hexameric motifs (red arrows).

We next probed the HP1γ RNAchIP for evidence of increased HP1γ RNA binding at differential splicing sites. Counting RNAchIP reads on the introns surrounding the differential splice sites revealed significantly higher levels of RNA in both input and IP samples (pVal=5×10^-6^ and 6×10^-10^, respectively), relative to a matching list of introns chosen randomly (Fig. 5d). This increase in read density was not observed when considering the host genes in their entirety (Fig. 5e). Together, these observations are indicative of HP1γ-RNA association having a local effect on splicing events.

Finally, as HP1γ also affects splicing of alternative exons (Fig. 5a and ref. ^25^), we examined RNAchIP read accumulation at exons annotated as “cassette exons” in the UCSC Alt Events database. Accumulation of reads at cassette exons was then compared to that observed either at introns or exons as defined in the RefSeq database. Consistent with HP1γ binding pre-mRNA, intronic reads were enriched in the IP fraction when compare to the input, while exonic reads were depleted (Fig. 5f). Interestingly, when restricting the analysis to cassette exons, we observed a moderate enrichment similar to the one seen on introns. Thus, when considering RNAchIP density profiles, cassette exons are intermediary between exons and introns. Next, to search for elements that could discriminate alternative exons from constant exons, we compared the densities of HP1γ binding motifs on cassette exons and RefSeq exons. We found that, while SINE repeats are excluded from RefSeq exons, as shown previously (Fig. 4e), cassette exons have a propensity to contain SINEs in their sequence (Fig. 5g). This suggests that cassette exon RNA is more prone to HP1γ binding than the average exonic RNA, a phenomenon that may contribute to the alternative usage of cassette exons. Collectively, our data suggest that CACACA motifs and the associated SINE repeats drive the targeting of HP1γ onto newly transcribed RNA, allowing HP1γ to participate in the accurate selection of genuine splice sites by the splicing machinery on chromatin (Fig. 5h).

## DISCUSSION

Elimination of intronic sequences from the pre-mRNA requires identification of the boundaries of exons with a resolution of one base. The initial recognition of a 5’ splice site (5’SS) relies on its base pairing with the U1 snRNP that then triggers the recruitment of the spliceosome. However, the consensus sequence recognized by the U1 snRNP is short and accepts many variants. Therefore, the mechanisms allowing U1 to distinguish between bona fide 5’SS and the many pseudo-5’SS present in the neighboring introns require the contribution of splicing co-regulators^35^. Here, we uncovered that HP1γ associates with repeated sequences present only in intronic RNA and that inactivation of the *HP1γ/Cbx3* gene results in increased usage of cryptic splice sites. These observations strongly suggest that HP1γ, by binding to intron-specific RNA sequences and tethering them to chromatin, participates in the positive selection of genuine splice sites. Interestingly, similar mechanism may apply at cassette exons that are intermediates between introns and exons in terms of content in sequences recognized by HP1γ.

Our strategy to identify chromatin-enriched RNA associated with HP1γ relied on a modification of our native chromatin immunoprecipitation assay. This approach revealed that the association of HP1γ with pre-messenger RNA on chromatin is based on its capacity to directly bind RNA molecules bearing preferentially a consensus CACACA or GAGAGA motif. The Hinge domain which contributes to RNA binding in HP1 proteins is not annotated in any of the common families of RNA binding domains. This Hinge region is an essentially unstructured peptidic segment between the two globular domains in HP1, and contains many positively charged basic residues, Lysines and Arginines, which create an electrostatic interface^7^. Considering the recent concept that heterochromatin-mediated gene silencing may occur in part through liquid-liquid phase separation involving association of HP1*α* with DNA^36, 37^, these physicochemical properties may be at play in the process of HP1γ association with RNA via its unstructured domain, creating local foci of chromatin condensation.

The two RNA consensus motifs we identified greatly enhanced the direct binding of purified HP1γ to RNA *in vitro*. However, an exact match with the consensus motifs was present in no more than half of the RNAchIP peaks. A possible reason for this discrepancy is that a subset of peaks contains motifs moderately divergent from the consensus. This suggests that HP1γ still recognizes an imperfect consensus, such as CACAC, the second motif present in the RNA probe used in our gel mobility shift assay. Interestingly, the CACACA motif is also recognized by hnRNP-L, an abundant nuclear protein that more generally binds intronic CA-repeats and CA-rich elements to regulate alternative splicing in mammals^38, 39^. We note however that genome-wide RNA binding sites of hnRNP-L do not significantly overlap with peaks of HP1γ-bound RNA (data not shown). Possibly, HP1γ and hnRNP-L may be competing for the same sites in the context of splicing regulation. The splicing factor 9G8 is also a potential competitor of HP1γ as it associates with the GAGAGA consensus motif^40^. Possibly, by recognizing multiple RNA motifs, HP1γ may act by interfering or cooperating with multiple positive and negative regulatory splicing factors, via their own RNA targeting motif within chromatin.

In humans, the short interspersed Alu repeats, when transcribed in antisense orientation generate a consensus 5’SS and therefore have contributed to the emergence of novel exon-exon junctions^41^. The SINE repeats involved in our model were in the sense orientation and were, in many of our examples, located near but not at differentially regulated exon-exon junctions. This strongly suggested that they were not used as a source of 5’SS. Instead, SINEs may function as anchors maintaining pre-messenger RNA on chromatin to facilitate the accurate co-transcriptional splicing. This would be compatible with our observations on the human CD44 gene where HP1γ contributes to regulation of alternative splicing while also tethering the pre-mRNA to the CD44 locus^25^. As a complement to this, we observed that a large majority of novel splicing junction involving cryptic splice sites in the absence of HP1γ were consistently spanning over shorter intronic regions than did similar splicing events in the presence of HP1γ. This raises the possibility that HP1γ also impacts on binding of splicing regulators by maintaining the pre-mRNA on chromatin. In this case, HP1γ may have an impact on the timely recruitment of regulatory subunits of the spliceosome, including the U2 snRNP that we found associated with HP1γ^30^. This configuration would provide a kinetic advantage of the strong splice donors and acceptors over cryptic sites, hence reinforcing the accuracy of splicing decisions.

Altogether, our results add a new key element to the model of chromatin-based regulation of pre-mRNA splicing by suggesting that HP1γ proteins may compete or synergize with splicing regulatory factors such as SR proteins and hnRNPs. In parallel, B4 SINE repeats, because they are enriched in CACACA motifs appear to be coincidentally good markers of the genome-wide distribution of HP1γ RNA-binding sites. In that sense, our data reconcile the heterochromatic and euchromatic functions of HP1 protein that are shown here to use similar repeat-binding properties both to silence interspersed repeats in intergenic region or inactive genes and to facilitate splicing of expressed genes. In fact, it seems that in the case of HP1γ, the function of heterochromatin-based silencing of SINE repeat elements within gene bodies has shifted towards regulation of alternative splicing at the locations of SINE repeats. Because of the diversity of HP1γ target genes and chromatin configurations, a wide spectrum of functional effects is observed, even from the specific point of view of splicing. The functional consequences of the CACACA-dependent targeting of HP1γ to RNA may have to be explored in specific cellular model systems for cell lineage specification.

## MATERIALS AND METHODS

### Cell lines and tissue culture

Mouse embryonic fibroblast (MEF)-derived cell lines were obtained exactly as previously described^16^. Briefly, HP1γ-expressing or KO cells were obtained by stable re-complementation of immortalized MEF HP1γ -/- cells by retroviral transduction using a retroviral vector carrying HP1γ cDNA, or an empty vector, respectively. Cells were grown in DMEM (Invitrogen) supplemented with 10% (v/v) fetal calf serum and 100 U.ml-1 penicillin-streptomycin. PMA-treated samples were obtained by addition of 100nM of Phorbol 12-Myristate 13-Acetate (PMA) in DMSO for 30 min on nearly confluent cells.

### Direct *in vitro* protein-RNA binding assay - Gel mobility shift assay

Assay was performed with bacterially expressed, purified GST fusion proteins, as depicted in Supplementary Fig. 2b, prepared as previously described^18^. Between 0.2 and 0.8 nmoles of GST-fusion proteins were incubated with 1 pmole of Cy3-labelled RNA oligonucleotide probes on ice for 20 min at 4°C in EMSA buffer (10mM Tris-HCl (pH 8.0), 50mM NaCl, 10% glycerol (w/v), 0.01% NP40, 0.1mg/ml BSA). The reaction was resolved by gel electrophoresis at 150 V for 20 min at +4°C, on a 5% native polyacrylamide gel (37.5:1) in 0.5x TBE buffer (45mM Tris-borate, 1mM EDTA). The gel was then immediately scanned on a Typhoon FLA 9000 (GE Healthcare), and subsequently stained with Coomassie blue R-250 (Sigma).

### Direct *in vitro* protein-RNA binding assay - North-western blot binding assay

Assay was performed essentially as described previously^12^. Briefly, bacterially expressed, purified GST-fusion proteins as above were separated by SDS-PAGE, transferred onto a nitrocellulose membrane (Biorad), renatured in PBS containing 5% of bovine serum albumin, and then hybridized for 1h at room temperature with biotinylated RNA probes that had been *in vitro* transcribed in the presence of biotin-16-UTP (Sigma) with T7 RNA polymerase (NEB) following the manufacturer’s instructions. After two washes and hybridization of Cy3-streptavidin (BioLegend, Inc.), membranes were scanned on a Typhoon FLA 9000 (GE Healthcare), and subsequently stained with Ponceau S (Sigma).

### Nuclei isolation and crosslinking

4×10^7^ cells treated or not with 100nM PMA for 30 min were washed directly on their tissue culture plate twice with ice cold Phosphate-Buffered Saline (PBS). All subsequent steps were performed at 4°C, unless otherwise specified. Cell were allowed to swell on ice for 5 min in 5ml of ice cold swelling buffer: (10mMTris-HCl pH7.5, 2mM MgCl2, 3mM CaCl2, supplemented before use with: 1x antiprotease (Roche), 0.5mM Na3VO4, 20mM *β*-Glycerophosphate, 80U/ml RNAsin (Promega), 0.1mM DTT). Cells were removed from the plate with a plastic cell scraper, transferred to a 15ml conical, and pelleted for 5 min at 4°C at 1600rpm. Cells were resuspended in 0.7ml of swelling buffer supplemented with 10% glycerol and 0.5% Igepal CA630 (Sigma), and gently pipetted up and down 15 times using a p1000 tip. Nuclei were pelleted for 5 min at 4°C at 2500rpm, and washed once in 1ml swelling buffer supplemented with 10% glycerol. Nuclei were then fixed with 0.3% formaldehyde in 300ul swelling buffer supplemented with 10% glycerol for 10 min at room temperature. Crosslinking was quenched during 10 min with 50mM glycine. Nuclei were pelleted for 5 min at 4°C at 3400rpm, and then washed once in 1ml swelling buffer.

### Chromatin-enriched RNA immunoprecipitation assay (RNAchIP)

Crosslinked nuclei were extracted in 0.8ml modified RIPA lysis buffer (50mM Tris-HCl pH7.5, 150mM NaCl, 0.5% sodium deoxycholate, 0.2% SDS, 1% NP-40, supplemented with 1x antiprotease (Roche), RNasin (Promega), and 0.5mM DTT). Cell suspension was sonicated using a Diagenode Bioruptor for 4 times 20s cycles at High amplitude. 7μl of Turbo DNase (Ambion, AM2238) and 7μl MgCl2 1M were added to sonicated material, incubated at 37°C for 10 min, and spun down at 12,000rpm for 10 min at 4°C. Ten percent of solubilized chromatin lysate was kept as input. The remaining volume was mixed with 350μl PBS and 200μl anti-FLAG magnetic beads suspension (Sigma, M8823) that were previously blocked in PBS containing 0,1% BSA, 0.5% Triton X-100 and 0.1% polyvinylpyrrolidone-40 (Sigma), and incubated at 4 °C for 2 h on a rotating wheel. Beads were then washed once in low salt wash buffer (1xPBS, 0.1% SDS, 0.5% NP-40), twice in high salt wash buffer (5x PBS, 0.1% SDS, 0.5% NP-40), and once more in low salt wash buffer. Beads were then eluted twice 10 min in 250μl elution buffer (150ng/μl 3xFLAG peptide (Sigma) in low salt wash buffer supplemented with RNasin). Eluates were combined for a total of 500μl. Eluates were then adjusted to 200mM NaCl and 10mM EDTA, and incubated with 10μg of Proteinase K at 50°C for 45 min, then placed at 65°C for 2h to reverse crosslinking. Samples were then subjected to phenol:chloroform extraction under acidic conditions followed with ethanol precipitation with Glycoblue (Ambion) as a carrier. Nucleic acid Pellets were then washed once in 75% ethanol, air-dried briefly, and resuspended in 30μl of RNase-free water for DNase treatment 20 min at 37°C, followed by RT-qPCR or library preparation.

### Reverse transcription and qPCR (RT-qPCR)

Reverse transcription was carried out with SuperScript III (Invitrogen) and random hexanucleotides for 1h at 50°C on 1μg RNA, quantified with a nanodrop (Thermo Scientific). Real-time quantitative PCR (qPCR) was carried out on a Stratagene Mx3005p with Brilliant III SYBR Green kits (Stratagene) according to the manufacturer’s instructions. Primer sequences are: Fos-in1-F, 5’-TGGAGACCACGAAGTGTTGGGAT-3’; Fos-in1-R, 5’-ATGGACACCTGCAACCTCTCAAGT-3’.

### Total RNA preparation and sequencing for transcriptome analysis

Total RNA was prepared out of HP1γ and KO cells by guanidinium thiocyanate-phenol-chloroform extraction according to the method of Chomczynski and Sacchi^42^, followed by proteinase K and DNAse treatments as described above. Total RNA library preparation and sequencing were performed on DNase-treated RNA samples by Novogene Co., Ltd, as a lncRNA sequencing service, including lncRNA stranded library preparation with rRNA depletion (Ribo-Zero Magnetic Kit), quantitation, pooling and PE 150 sequencing (30G raw data-100M raw reads/sample) on Illumina HiSeq 2500 platform.

### RNAchIP Library preparation

RNA quality and yield were assessed by the RNA integrity number (RIN) algorithm, using the 2100 Bioanalyzer. Directional libraries were prepared using the Smarter Stranded Total RNA-Seq kit-Pico Input Mammalian kit following the manufacturer’s instructions (Clontech, 635005). The quality of all libraries was verified with the DNA-1000 kit (Agilent) on a 2100 Bioanalyzer and quantification was performed with Quant-It assays on a Qubit 3.0 fluorometer (Invitrogen). Clusters were generated for the resulting libraries, with the Illumina HiSeq SR Cluster Kit v4 reagents. Sequencing was performed using the Illumina HiSeq 2500 system and HiSeq SBS kit v4 reagents. Runs were carried out over 65 cycles, including seven indexing cycles, to obtain 65-bp single-end reads. Sequencing data were then processed with the Illumina Pipeline software, Casava (v.1.9).

### Bioinformatics analysis

Bioinformatics analysis of the RNAchIP-seq was performed using the RNA-seq pipeline from Sequana^43^. Reads were cleaned of adapter sequences and low-quality sequences using cutadapt (v.1.11)^44^. Only sequences at least 25 nt in length were considered for further analysis. STAR (v.2.5.0a)^45^ (parameters: --outFilterMultimapNmax 30 --outSAMmultNmax 1--outMultimapperOrder Random) was used for alignment on the reference genome (Mus musculus mm9 from UCSC). Genes were counted using featureCounts (v.1.4.6-p3)^46^ from Subreads package (parameters: -t CDS -g ID -s 1). MACS2 (v.2.1.0)^47^ was used to call HP1-binding peaks on RNA-ChIP data (parameters: --nomodel --extsize=150 -q 0.1). Bamcoverage from Deeptools^48^ was used to produce normalized BigWig files to 1X. Finally, Bedtools (v.2.25.0)^49^ closestBed (parameters: -D ref -mdb each) was used to annotate each peak from all conditions with related and public ChIP-seq data from Gene Expression Omnibus database (H3k4me3, RNA pol II, H3K9me3, H3K27me3; GEO sample accessions: GSM769029, GSM918761, GSM2339533, GSM946547, respectively). All random controls were performed with Bedtools (v.2.25.0) Shuffle over intragenic intervals (parameter: -incl option to restrict randomizations to within gene bodies). Average profiles were obtained using plotProfile (parameter: --perGroup) and heatmaps using plotHeatmap (default parameters) out of matrices generated by computeMatrix (parameters: --referencePoint center, Fig. 1-3, or scale-regions, Fig.4, 5), from Deeptools, on BigWig files of RNA-ChIP reads or intragenic SINE, LINE, or hexameric CACACA or GAGAGA motifs. SINE and LINE repeats were obtained from the RepeatMasker database at UCSC. CACACA or GAGAGA strict hexameric motifs were obtained with the scanMotifGenomeWide.pl tool from the Homer package (v. 4.9.1). Intragenic features were obtained with intersectBed from Bedtools, with strand orientation based on the Mus_musculus mm9 reference database (Ensembl release 67), and then converted into BigWig files for analysis via computeMatrix. All genome views were done using the Integrative Genomics Viewer software (IGV)^50^.

### Statistical analysis

Each count dataset was analyzed using R (v.3.4.1) and the Bioconductor package DESeq2 (v.1.16.0)^51^ to test for the differential gene expression (DGE). The normalization and dispersion estimation were performed with DESeq2 using the default parameters and statistical tests for differential expression were performed applying the independent filtering algorithm. A generalized linear model was set in order to test for the differential expression between the biological conditions. For RNAchIP-seq, triplicates IP samples were compared to input samples in each PMA-treated and unstimulated samples. For RNA-seq, triplicate HP1 samples were compared to KO samples in each PMA-treated and unstimulated cells. For each pairwise comparison, raw p-values were adjusted for multiple testing according to the Benjamini and Hochberg (BH) procedure^52^ and genes with an adjusted P-value lower than 0.05 were considered differentially expressed.

**Motif discovery by RSAT** was performed online on the RSAT peak motif search interface (http://rsat.sb-roscoff.fr/)^53^. First, sequences of the merged RNAchIPseq peaks from PMA-treated and unstimulated conditions were obtained in fasta format with strand orientation based on the Mus_musculus mm9 reference database (Ensembl release 67). The fasta files were then used as queries for RSAT peak-motif disccovery (parameters: peak-motifs -v 1 - markov auto -disco oligos, positions -nmotifs 5 -minol 6 -maxol 8 -no_merge_lengths −1str - origin center). The e-value associated with each discovered motif represents the expected number of patterns which would be returned at random for a given probability of occurrence (P-value).

### Analysis of differential splicing events in the transcriptome by MAJIQ

Alternative splicing events occuring in the transcriptome between different conditions were analyzed by the MAJIQ computational framework (v.2.1-179b437)^34^ with default parameters. For this purpose, the transcriptome was aligned with STAR (parameters: -- outFilterMismatchNmax 1 --outMultimapperOrder Random --outSAMmultNmax 1 -- outFilterMultimapNmax 30) on the mouse GRCm38/mm10 unmasked genome. The definition of annotated and *de novo* junctions were based on the mouse GRCm38/mm10 Ensembl annotation database. Analyses were performed by pairwise comparisons between experimental conditions, namely HP1γ versus KO, and HP1γ +PMA versus KO+PMA, for each triplicate. Local splicing variations (LSVs) were detected by the software and their relative abundance (PSI) quantified for each condition, leading to a relative change (dPSI) between HP1γ and KO for each junction involved in the LSV. The default threshold of change of |dPSI| >= 0.2 (20%) between conditions was used. Genomic locations of LSVs were converted from mm10 to mm9 by the Liftover tool at UCSC genome browser before further analysis.

### Data availability

RNAchIP-seq and RNA-seq data have been deposited in the NCBI Gene Expression Omnibus database under GEO accession number GSE133267.

## ACKNOWLEDGEMENTS

We are grateful to all members of the Epigenetic Regulation unit for helpful discussions, Madeleine Moscatelli and Cynthia Bezier for helpful preliminary experiments, Catherine Bodin for technical assistance and Edith Ollivier for administrative assistance. Thanks to Caroline Proux for her expertise in library preparation and Illumina sequencing, and to Eric Batsché for critically reading the manuscript.

This work was supported by Institut National de la Santé et la Recherche Médicale (Inserm; C.R.), Centre National de la Recherche Scientifique (CNRS; C.M.), with grants from Agence Nationale de la Recherche (ANR-11-BSV8–0013) and REVIVE––Investissement d’Avenir (to E.K. and C.M.). J.Y. is part of the Pasteur - Paris University (PPU) International PhD Program. This program has received funding from the European Union’s Horizon 2020 research and innovation programme under the Marie Sklodowska-Curie grant agreement No 665807.

## AUTHOR CONTRIBUTIONS

C.R. conceived, carried out and analyzed the experiments and performed some bioinformatics analyses. R.L. performed bioinformatics analyses for the RNAchIPseq and transcriptome. M.C. performed bioinformatics analyses with MAJIQ. H.V. performed statistical analyses. J.Y. carried out the RNA preparations for the transcriptome. E.K. contributed to bioinformatics analysis of RNAchIP peak colocalizations. C.R. and C.M. contributed to the overall orientations of the project. C.R. wrote the manuscript and all authors were involved in revising it critically for important intellectual content.

## COMPETING INTERESTS

Authors declare no competing interests.

