## Supplementary Material for "HP1γ binding pre-mRNA at intronic repeats increases splicing fidelity and regulates alternative exon usage"

**Supplementary Fig. 1**

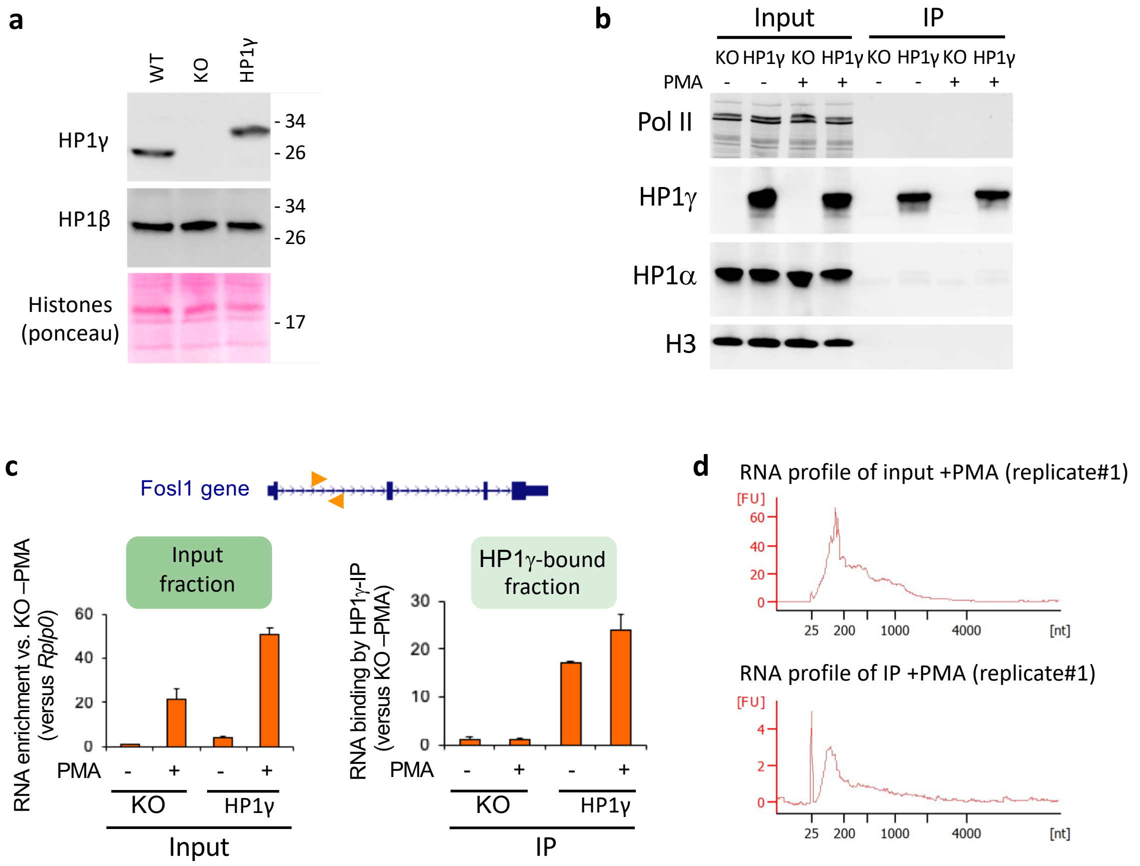


**Supplementary Fig. 1** **a**, Top panels, western blot analysis of HP1γ and HP1β protein levels in MEF-derived cells expressing or not FLAG-tagged HP1γ (HP1γ or KO, respectively), compared to WT MEF-derived cells. Bottom panel, total protein staining by Ponceau S. Panel is a section of the blot centered on 17kD, showing Histones. **b**, Western blot analysis of the fractions used in the RNAchIP strategy in the conditions tested in the HP1γ-expressing cells (HP1γ), compared to HP1γ -/- cells (KO). **c**, Relative quantities of RNA using primer pairs aligning in the intron1 of the *Fosl1* gene (orange arrowheads), detected by RT-qPCR on the RNAchIP samples. Graphes represent mean and s.d., *n*=3 independent experiments. **d**, Virtual gel profiles showing the size range of RNA fragments in both input and IP samples from HP1γ cells, obtained by Bioanalyzer (Agilent).

### Supplementary Fig. 2

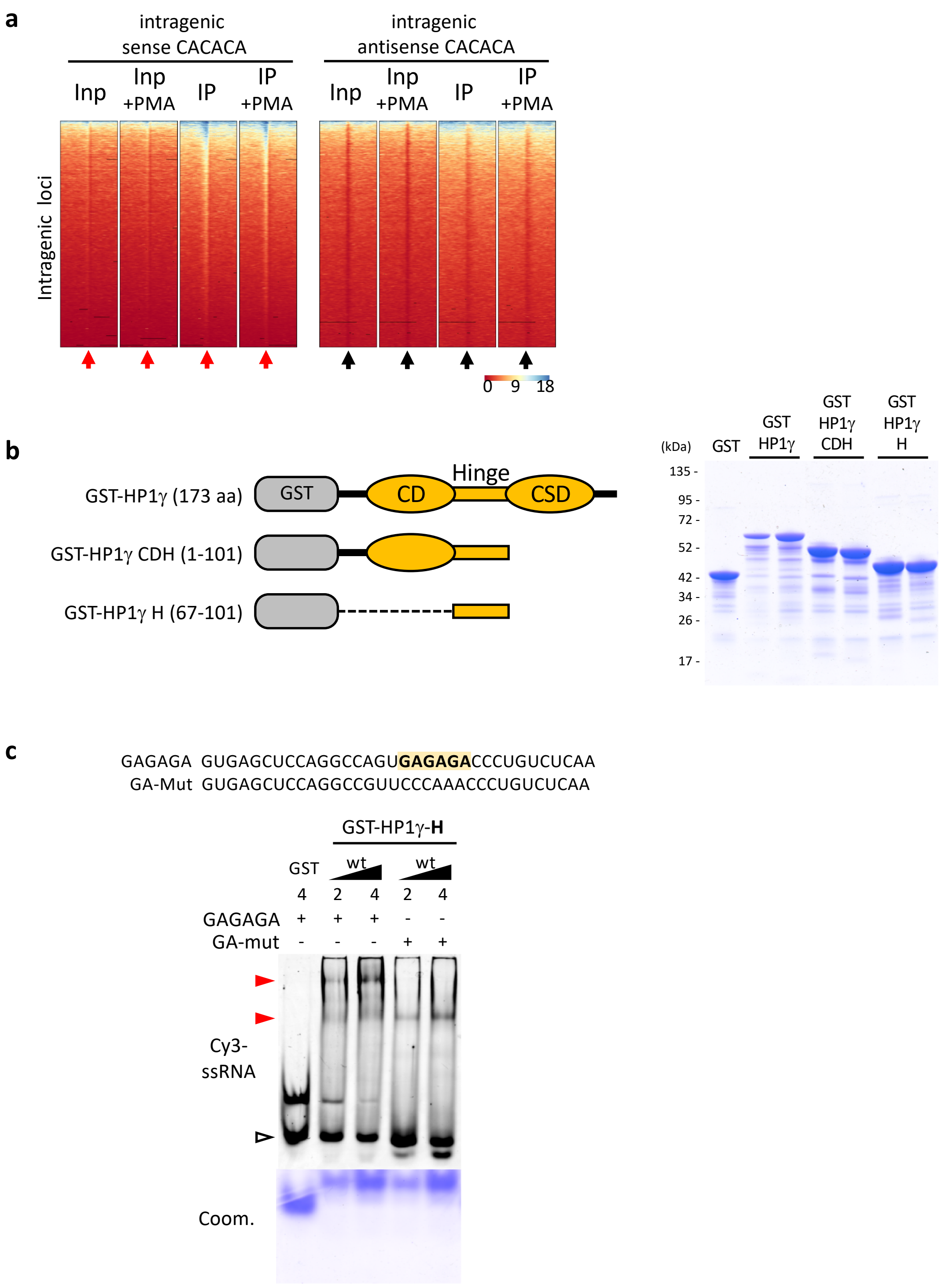


**Supplementary Fig. 2** **a**, Heat maps of IP and input signal in one triplicate sample for each condition over ±2kb centered (arrows) on intragenic CACACA hexameric motifs, corresponding to the average profiles depicted in Fig. **2b**. **b**, Left, schematic representation of the GST-HP1γ constructions used in gel mobility shift assay, depicting the chromo- and chromoshadow- globular domains, as well as the unstructured Hinge domain (CD, CSD, Hinge, respectively). Right, Coomassie blue staining of duplicate samples of the bacterially expressed, purified GST- HP1γ fusion proteins. **c**, Gel mobility shift assay performed in the same conditions as in **Fig. 2d**, on a GAGAGA-containing RNA oligonucleotide probe (GAGAGA) and on a control mutated motif-containing RNA probe (GA-mut).

**Supplementary Fig. 3**

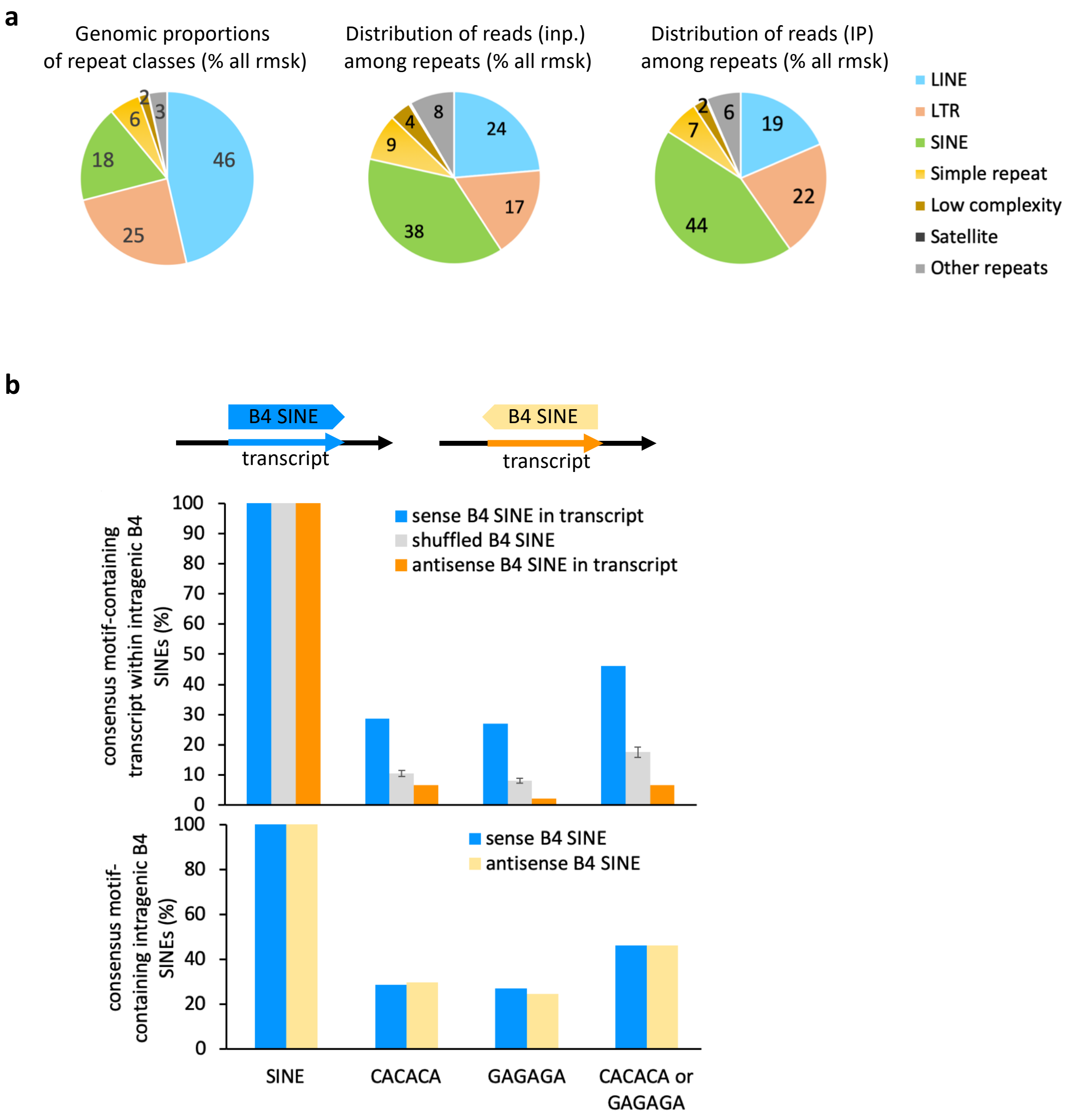


**Supplementary Fig. 3 a,** Pie charts depicting the proportions of repeat classes of the Repeat Masker database, in terms of their cumulated genomic size (left chart), percentage of RNA sequencing reads from input (middle chart), and IP (right chart) counted per repeat classes as a percentage of all repeats, on the basis of all uniquely aligned reads. **b,** Intragenic B4 SINE repeats have a CACACA or GAGAGA motif density which is linked to their orientation in the gene, not with the proportions of CACACA in their intrinsic sequence. Top, schematic representation of the features. Middle, proportions of intragenic B4 SINE repeats which contain the depicted hexameric motifs when the sequence is read in the same orientation as the overlapping gene. Bottom, both sense and antisense B4 SINE repeats contain identical intrinsic motif density in their sequence. Graphes represent mean and s.d. for the randomized intervals, *n*=3.

**Supplementary Fig. 4**

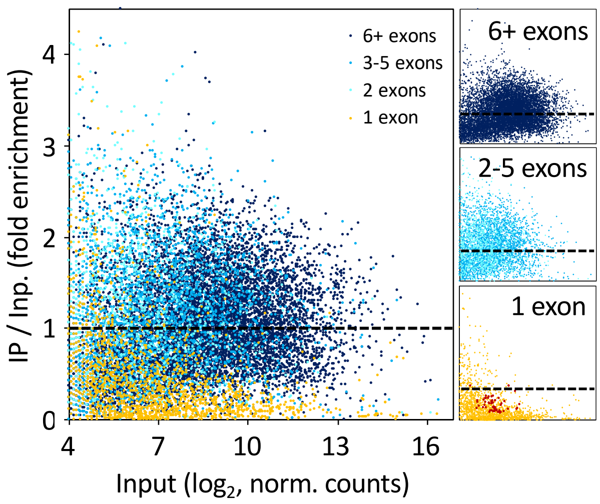


**Supplementary Fig. 4** Scatter plot of fold enrichment and input represented as in c, corresponding to Fig. 4a. Genes were color-coded according to their number of exons (Orange, 1 exon; light to dark blue, 2 to 6 and more exons, respectively), and sorted in individual panels by color. Red dots highlight intron-less histone genes.

**Supplementary Fig. 5**

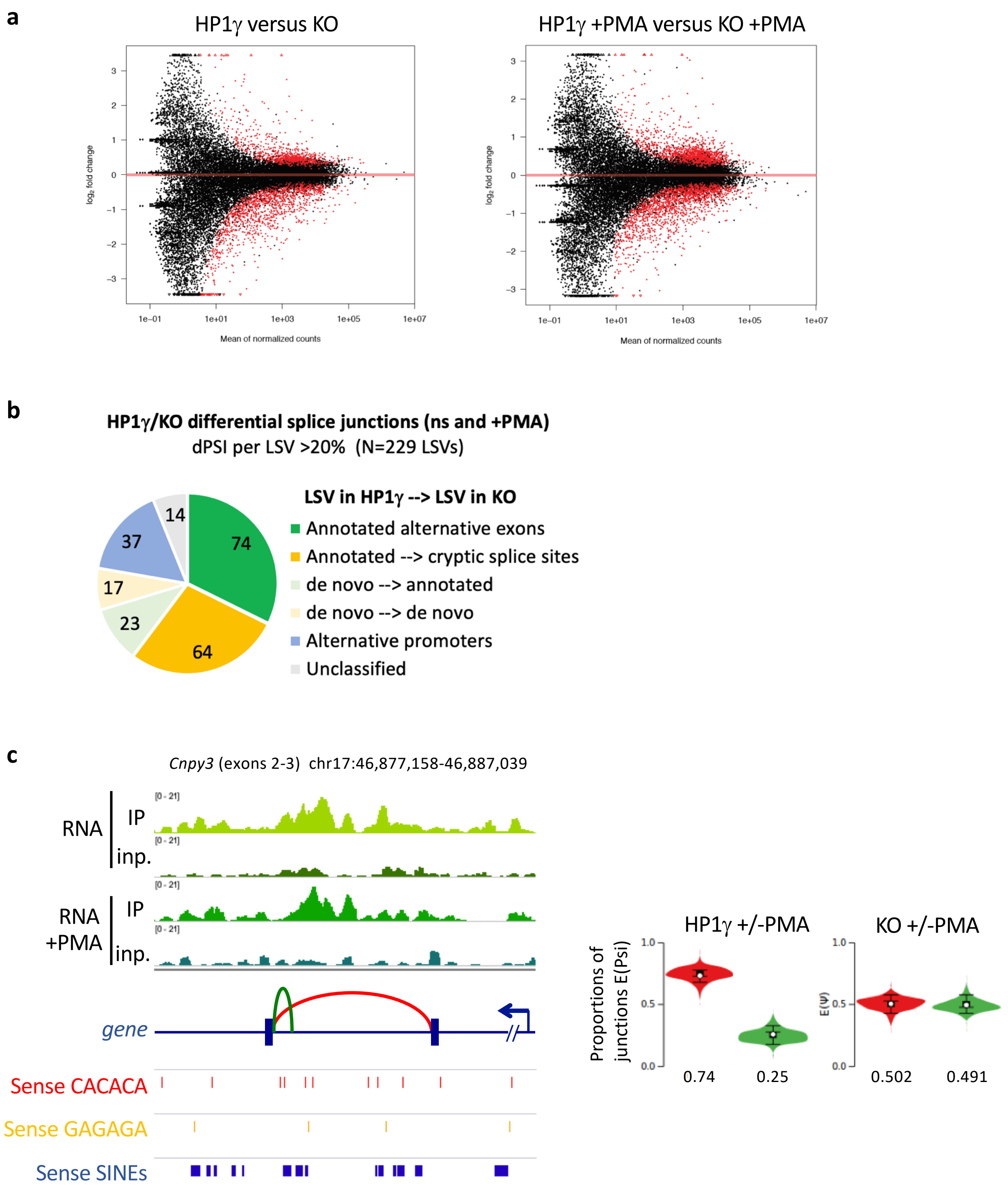


**Supplementary Fig. 5** Differential splicing events in HP1γ and KO transcriptomes. **a,** MA-plot of the transcriptome dataset, *n*=3 biological replicates. Red dots represent significantly differentially expressed features, as specified in the methods section. **b,** Number of high confidence LSVs d(PSI)>=0.2 between HP1γ and KO. Unclassified are Junctions in rRNAs, satellite repeats, or undefined junction changes (not matching any gene). **c,** Left, genome view showing the *de novo* cryptic splicing event depicted in Fig **5b**, together with RNAchIP read density profiles (top), and position of the depicted oriented repeated motifs (bottom). Right, plot showing quantifications of expected abundance computed by MAJIQ, as in Fig. **5b.**

Full-size images related to **Supplementary Fig. 1b.**

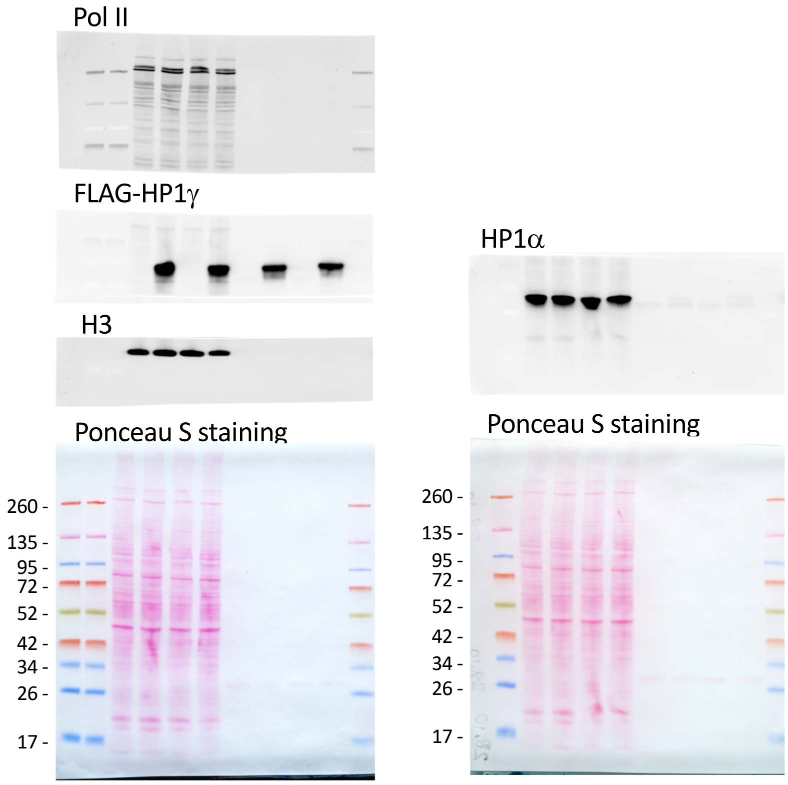


Full size images related to **Fig. 3h**

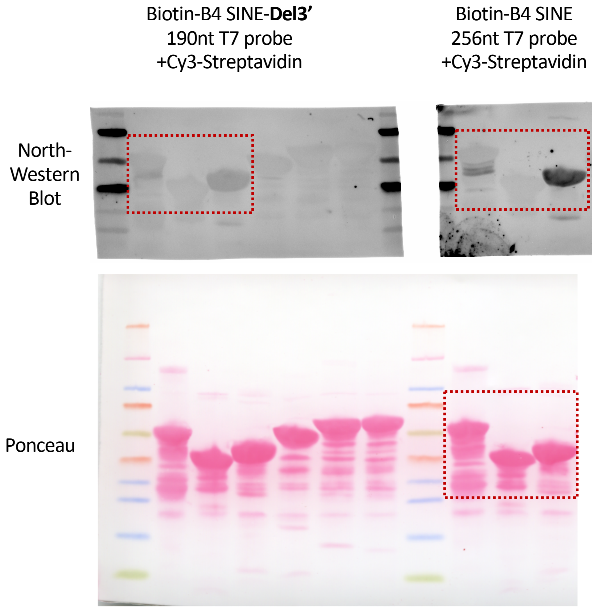
